# COPILOT: a Containerised wOrkflow for Processing ILlumina genOtyping daTa

**DOI:** 10.1101/2021.07.26.453753

**Authors:** Hamel Patel, Sang-hyuck Lee, Gerome Breen, Stephen Menzel, Oyesola Ojewunmi, Richard J.B Dobson

**Affiliations:** Department of Biostatistics and Health Informatics, Institute of Psychiatry, Psychology and Neuroscience (IoPPN), King’s College London, 16 De Crespigny Park, London, SE5 8AF, UK; NIHR Biomedical Research Centre at South London and Maudsley NHS Foundation Trust and King’s College London, London, UK; Social, Genetic and Developmental Psychiatry Centre, Institute of Psychiatry, Psychology and Neuroscience, King’s College London, London, UK; Comprehensive Cancer Centre, School of Cancer and Pharmaceutical Sciences, King’s College London; Health Data Research UK London, University College London, London, UK; Institute of Health Informatics, University College London, London, UK; NIHR Biomedical Research Centre at University College London Hospitals NHS Foundation Trust, London, UK

**Keywords:** Illumina, microarrays, genotype, container, pipeline, QC, GWAS, GenomeStudio

## Abstract

**Background:** The Illumina genotyping microarrays generate data in image format, which is processed by the platform-specific software GenomeStudio, followed by an array of complex bioinformatics analyses. This process can be time-consuming, lead to reproducibility errors, and be a daunting task for novice bioinformaticians.

**Results:** Here we introduce the COPILOT (**C**ontainerised w**O**rkflow for **P**rocessing **IL**lumina gen**O**typing da**T**a) protocol, which provides an in-depth and clear guide to process raw Illumina genotype data in GenomeStudio, followed by a containerised workflow to automate an array of complex bioinformatics analyses involved in a GWAS quality control (QC). The COPILOT protocol was applied to two independent cohorts consisting of 2791 and 479 samples genotyped on the Infinium Global Screening (GSA) array with Multi-disease (MD) drop-in (~750,000 markers) and the Infinium H3Africa consortium array (~2,200,000 markers) respectively. Following the COPILOT protocol, an average sample quality improvement of 1.24% was observed across sample call rates, with notable improvement for low-quality samples. For example, from the 3270 samples processed, 141 samples had an initial sample call rate below 98%, averaging 96.6% (95% CI 95.6-97.7%), which is considered below the acceptable sample call rate threshold for a typical GWAS analysis. However, following the COPILOT protocol, all 141 samples had a call rate above 98% after QC and averaged 99.6% (95% CI 99.5-99.7%). In addition, the COPILOT pipeline automatically identified potential data issues, including gender discrepancies, heterozygosity outliers, related individuals, and population outliers through ancestry estimation.

**Conclusions:** The COPILOT protocol makes processing Illumina genotyping data transparent, effortless and reproducible. The container is deployable on multiple platforms, improves data quality, and the end product is analysis-ready PLINK formatted data, with a comprehensive and interactive summary report to guide the user for further data analyses.

## Background

A genome-wide association study (GWAS) is an approach to identify genetic variants associated with a particular disease or phenotypic trait. Microarray-based GWAS, such as Illumina genotyping, remains a common approach for identifying these genetic associations across the whole genome. The Illumina genotyping microarrays generate data in image form, which is initially processed using the platform-specific software, GenomeStudio. The data is then further processed by a series of complex bioinformatics analyses, which depend on various pieces of software, programming languages and their dependencies to be installed and configured correctly. This can be a time-consuming process and can be a daunting task for bioinformaticians/statistical geneticists. Furthermore, the relatively easy installation of software is vital to encourage reproducible data analysis (1).

Here we introduce COPILOT, a **C**ontainerised w**O**rkflow for **P**rocessing **IL**lumina gen**O**typing da**T**a. The automated and easy to install pipeline consists of a series of bash, C/C++, R, and Python programs containerised using the docker framework, ready to execute on multiple operating platforms with minimal effort from the user. The pipeline will take the output from GenomeStudio, apply a secondary clustering algorithm to improve call rates (2) and apply multiple analyses for QC, including identification of any genotypic and phenotypic gender discrepancies, calculation of Identity-by-descent (IBD), perform heterozygosity testing and estimate sample ancestry based on the 1000 genome reference panel. The processed data is output in PLINK format and is accompanied by a detailed interactive summary report with informative plots and explanations to aid the user for downstream analyses.

## Implementation

### Architecture and implementation

COPILOT has been compiled into a docker image, which can house complex analysis pipelines consisting of several configured software and can be deployed on multiple operating platforms (3). Comprehensive documentation, including downloading and installing COPILOT, is available at https://khp-informatics.github.io/COPILOT/COPILOT_user_guide.html. The documentation includes a working tutorial with real-world genotyping data and an example of the resulting interactive summary report. First, the raw genotyping intensity data must be processed using the Illumina GenomeStudio software using guidelines previously published (4–6) or following our comprehensive GenomeStudio QC protocol, which contains additional QC procedures and visual aids to guide the user. Our GenomeStudio QC protocol can be accessed at https://khp-informatics.github.io/COPILOT/GenomeStudio_genotyping_SOP.html. We collectively refer to our comprehensive GenomeStudio QC guidelines and the COPILOT container as the COPILOT protocol.

### Data input and output

The COPILOT container requires three input files. The first is a GenomeStudio report file containing the processed genotype intensity values, the second is the genotyping manifest file containing information on the probe design, and the third is a gender file containing phenotypic gender. The COPILOT pipeline is executed using an execution script that specifies the input files, the docker image, and the desired output location for all the generated files. Details on generating the input files and executing the container are available in the online documentation. The pipeline outputs the processed data in PLINK format, ready for further analysis and a comprehensive summary report.

### Data processing

The COPILOT container starts by performing pre-analysis checks of the input data, calculates basic statistics, and prepares for the zCall rare variant calling algorithm. The zCall caller is an established software (2), which attempts to assign genotype calls to SNPs that have been missed by the GenomeStudio GenCall algorithm, which is commonly the rare variants. This process essentially increases the overall sample call rates, which effectively increases data quality. COPILOT then converts data to PLINK format and uses the manifest file to update all alleles in AB format to Illumina TOP strand and then performs multiple analyses as recommended in (7). This includes iteratively removing SNPs and samples to a user-defined threshold, pruning data for linkage disequilibrium (LD) and removing high regions of LD and non-autosomal regions, identify potential sex discrepancies, calculating Identity-by-descent (IBD), performing a heterozygosity test and calculating ancestry of each sample based on the 1000 genome reference panel (8).

Genotyping data often consists of duplicate sample ID’s, which can cause issues during data processing. COPILOT handles duplicate sample IDs by assigning temporary unique sample ID’s to duplicate samples, allowing each duplicate sample to be independently treated during the data processing stage. The resulting processed data is provided back to the user in PLINK format, with any duplicate sample ID’s reverted to the original ID, but a record kept for users to re-identify them if required. An outline of the COPILOT process is provided in Figure 1.

**Figure 1:**
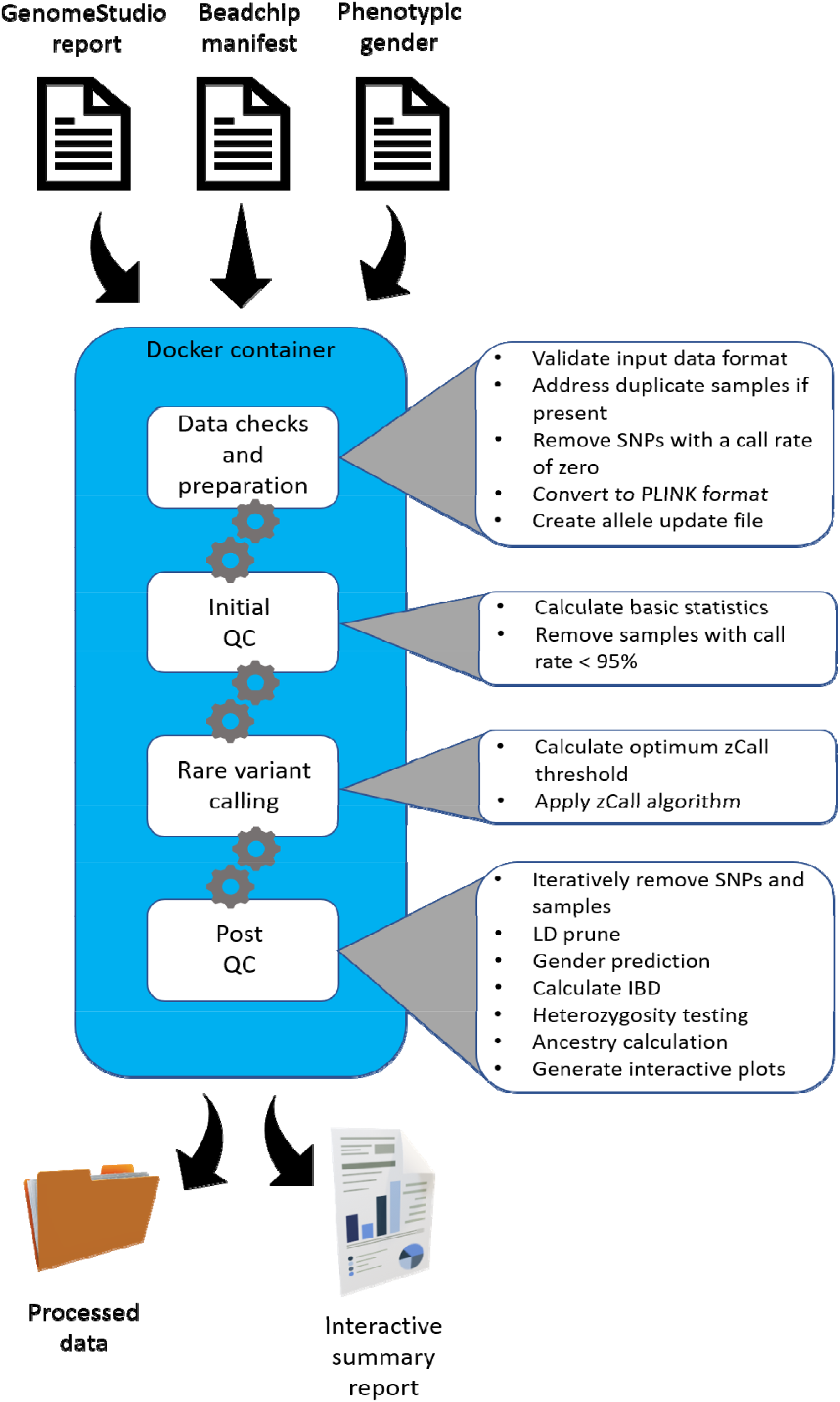
Outline of COPILOT container process.

## Results

### Performance

The COPILOT protocol is currently deployed in the KING’S College London’s (KCL) Institute of Psychiatry, Psychology and Neuroscience’s (IoPPN) Genomics & Biomarker Core Facility, and has been successfully used to process thousands of samples on various genotyping chips, including the Infinium Core, Exome, CoreExome, OmniExpress, Omni2.5, Omni5, PsychArray, Multi-Ethnic, Global Screening Array range and H3Africa consortium arrays. The container takes approximately 60 minutes to process ~300 samples on the Infinium HumanCoreExome Array (~550,000 SNPs) using a machine with six dual cores and 32GB ram. A substantially larger project with ~500 samples on the H3Africa Consortium Array (~2.3 million SNPs) takes approximately 30 hours to process.

### Sample quality improvement – case study

The sample call rate is the fraction of the SNPs with a genotype call for a given sample, with higher sample call rates indicative of better sample quality. The COPILOT protocol was applied to 3270 samples from two different cohorts using two different Illumina genotyping arrays. The first cohort is a mental health disorder consisting of 2791 samples collected as either blood, buccal swab or saliva, and genotyped using the Infinium Global Screening (GSA) array with Multi-disease (MD) drop-in (~750,000 markers). The second cohort is a sickle cell cohort consisting of 479 samples collected as blood samples and genotyped using the Infinium H3Africa consortium array (~2,200,000 markers).

The cohorts were independently processed in GenomeStudio using the default GenCall clustering algorithm with extreme outliers removed (samples with a call rate below 90%). An initial average sample call rate of 98.55% (95% CI 98.58-98.67%) was achieved for the mental health cohort, and 99.19% (95% CI 99.14-99.22%) was achieved for the sickle cell cohorts before any QC. Following the COPILOT QC protocol (including GenomeStudio QC and COPILOT container) and without requiring removal of additional samples, the average sample call rates improved to 99.86% (95% CI 99.86-99.89) and 99.93 (95% CI 99.93-99.95) respectively, averaging an improvement of 1.24% across the 3270 samples. Notably, the sample call rates significantly improved for samples at the lower end of the sample quality spectrum (Figure 2). The sample call rate threshold used to exclude samples from a typical GWAS varies, with 98% commonly used (7). The mental health and sickle cell cohort consisted of 129 and 12 samples, respectively, where sample call rates were below 98%. The average call rates for these 141 samples were 96.6% (95% CI 95.6-97.7%) before QC; however, following the COPILOT protocol, these 141 samples all attained a call rate above 98% and averaged 99.6% (95% CI 99.5-99.7%).

**Figure 2:**
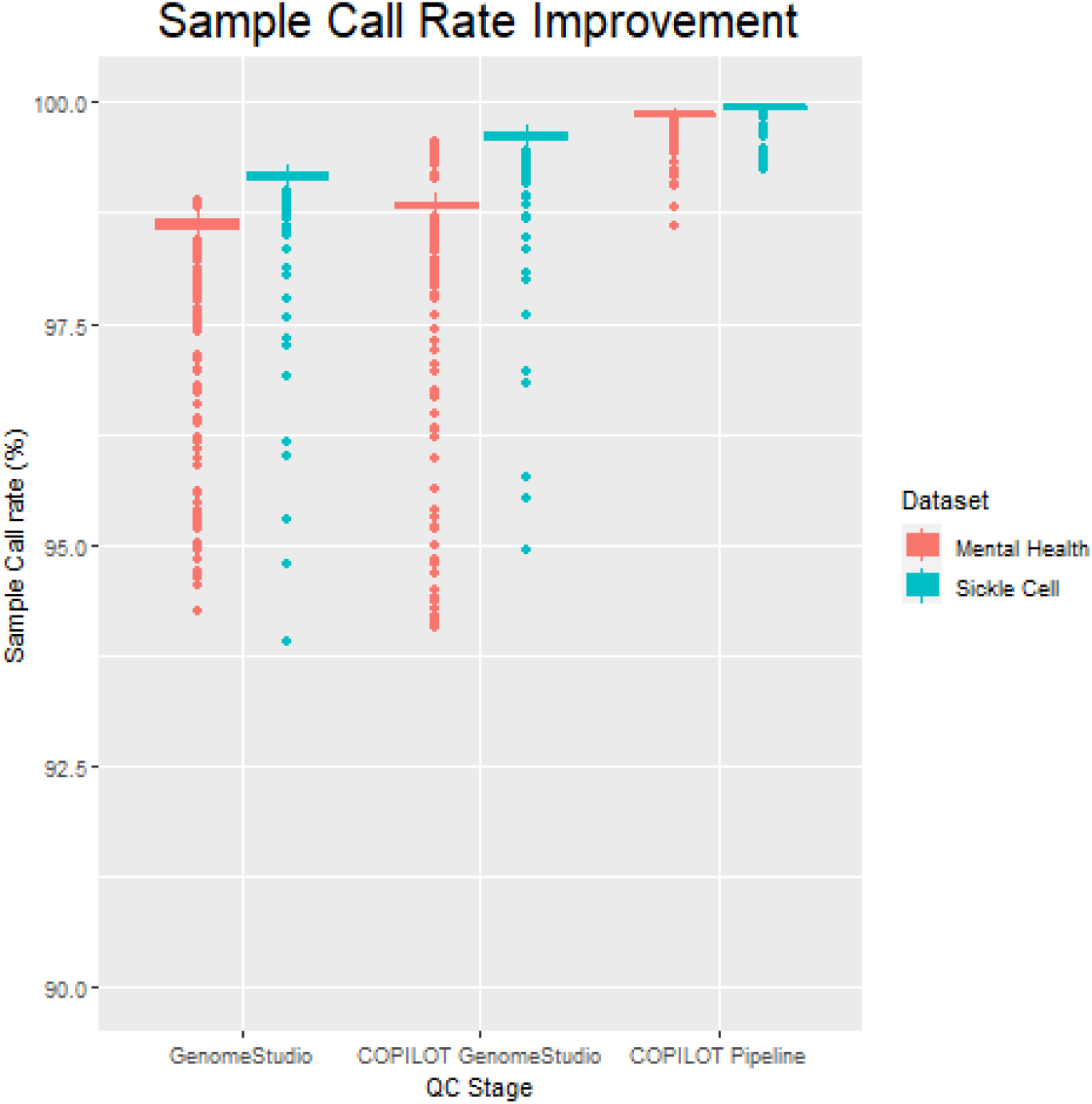
Sample quality improvement observed on two cohorts through three QC stages of the COPILOT protocol. The first cohort is a mental health disorder consisting of 2791 samples collected as either blood, buccal swab or saliva, and genotyped using the Infinium Global Screening (GSA) array with Multi-disease (MD) drop-in (~750,000 markers). The second is a sickle cell cohort consisting of 479 samples collected as blood samples and genotyped using the Infinium H3Africa consortium array (~2,200,000 markers). The QC stages are 1) “GenomeStudio”, where samples are clustered using the GenCall algorithm, 2) “COPILOT GenomeStudio”, where samples are processed in GenomeStudio using the comprehensive COPILOT GenomeStudio QC protocol, and 3) “COPILOTpipeline”, where samples are processed through the containerised COPILOT pipeline. Samples that failed clustering in the GenomeStudio stage have been excluded from this plot to make sample improvement comparable.

### Automated identification of problematic samples – case study

The COPILOT container automates the identification of potentially problematic samples for a typical GWAS and provides findings through an informative and interactive summary report (example provided here at https://khp-informatics.github.io/COPILOT/README_summary_report.html). For example, COPILOT was used to process the sickle cell cohort consisting of 479 patients. As a result, COPILOT identified genotypic and phenotypic gender discrepancies (Figure 3), heterozygosity outliers (Figure 4), related samples (Figure 5), sample ancestry based on the 1000 genome reference panel (Figure 6), and provide an overall summary of potential problematic samples (Figure 7).

**Figure 3:**
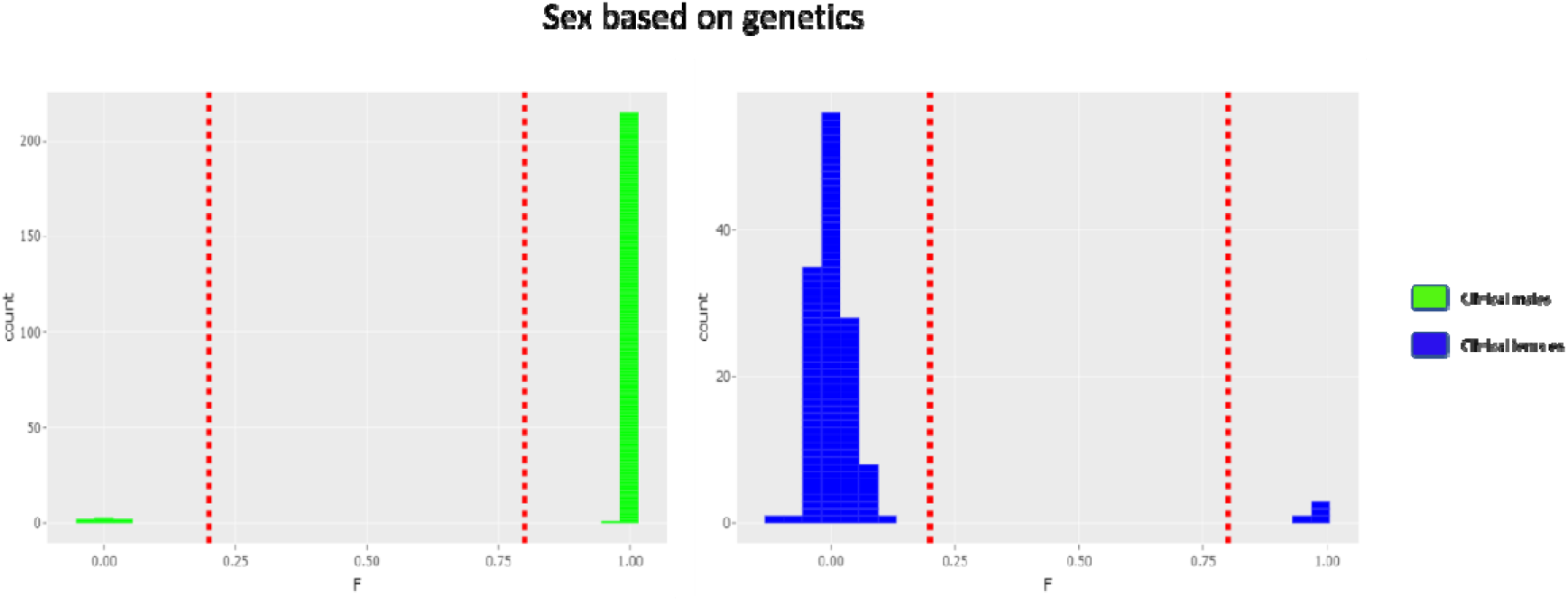
Gender discrepancies in the sickle cell cohort. The F-statistics is calculated for all samples and represents the expected level of heterozygosity on the X-chromosome. Samples with an F statistic above 0.8 are presumed to be genetically “Male”, while samples with an F statistic below 0.2 are presumed to be genetically “Female”. COPILOT identified 13 gender discrepancies in the Sickle cell cohort where the clinically defined gender differed from genetics.

**Figure 4:**
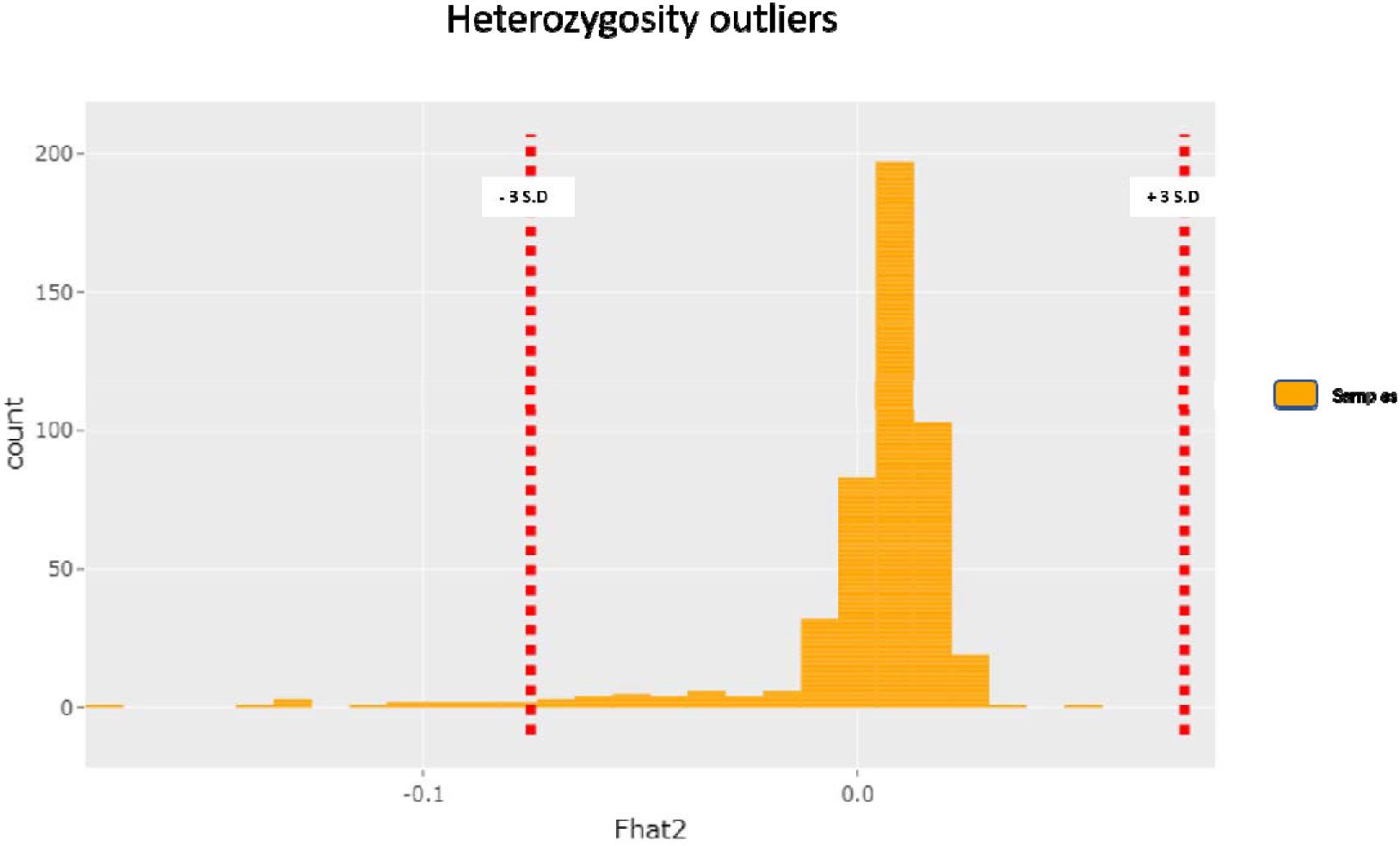
Heterozygosity testing in the sickle cell cohort. Samples that deviate from the expected heterozygosity, compared to the overall heterozygosity rate of the study, can aid in identifying problematic samples. High levels of heterozygosity can indicate low-quality samples, while low levels of heterozygosity can be due to inbreeding. COPILOT identified 14 heterozygosity outliers in the Sickle cell cohort.

**Figure 5:**
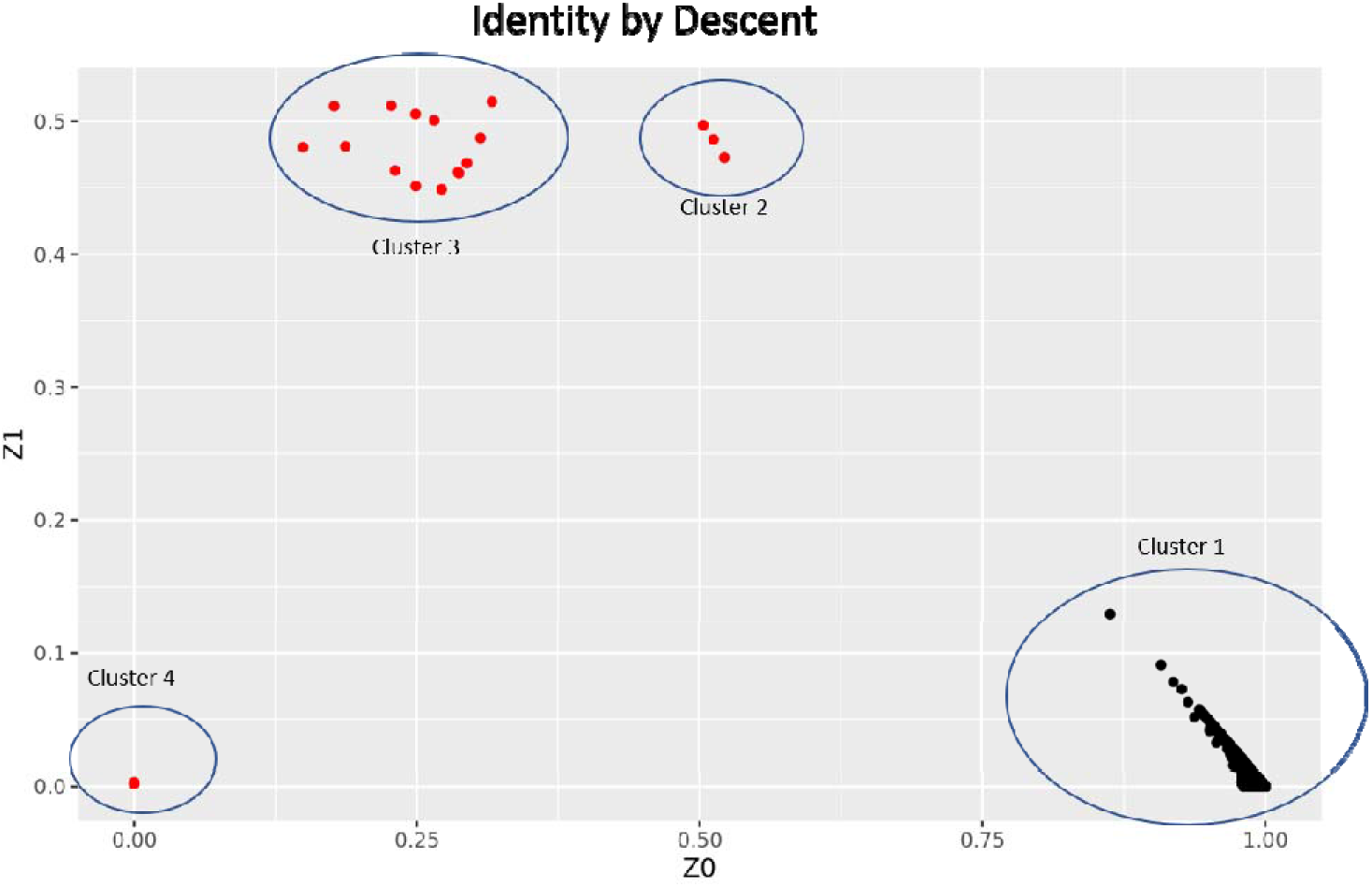
Identity-by-Descent calculation in the sickle cell cohort. A typical GWAS assumes all subjects are unrelated; therefore, closely related samples can lead to biased errors in SNP effects if not correctly addressed. If self-reported relationship information is available, IBD can identify potential sample mix-ups and/or cross-sample contamination. Individuals with an IBD pi-hat metric over 0.1875 (halfway between a second and third-degree relative) have been identified as related. The z0, z1 and z2 metrics indicate the proportion of the same copies of alleles (z0 = 0 copies, z1= 1 copy and z2 = 2 copies) shared between two individuals, with PI_HAT calculated as P(IBD=2) + 0.5*P(IBD=1). These metrics can be used to estimate the type of relation. Analysing the sickle cell cohort identified four clusters where Cluster 1 (pi-hat < 0.1875) indicates unrelated samples, cluster 2 (z0 = 0.5 and z1 = 0.5) indicates half-siblings, cluster 3 (z0 = 0.25 and z1 = 0.5) indicates full siblings and cluster 4 (z0 = 0 and z1 = 0) indicates potential contamination or duplicate samples.

**Figure 6:**
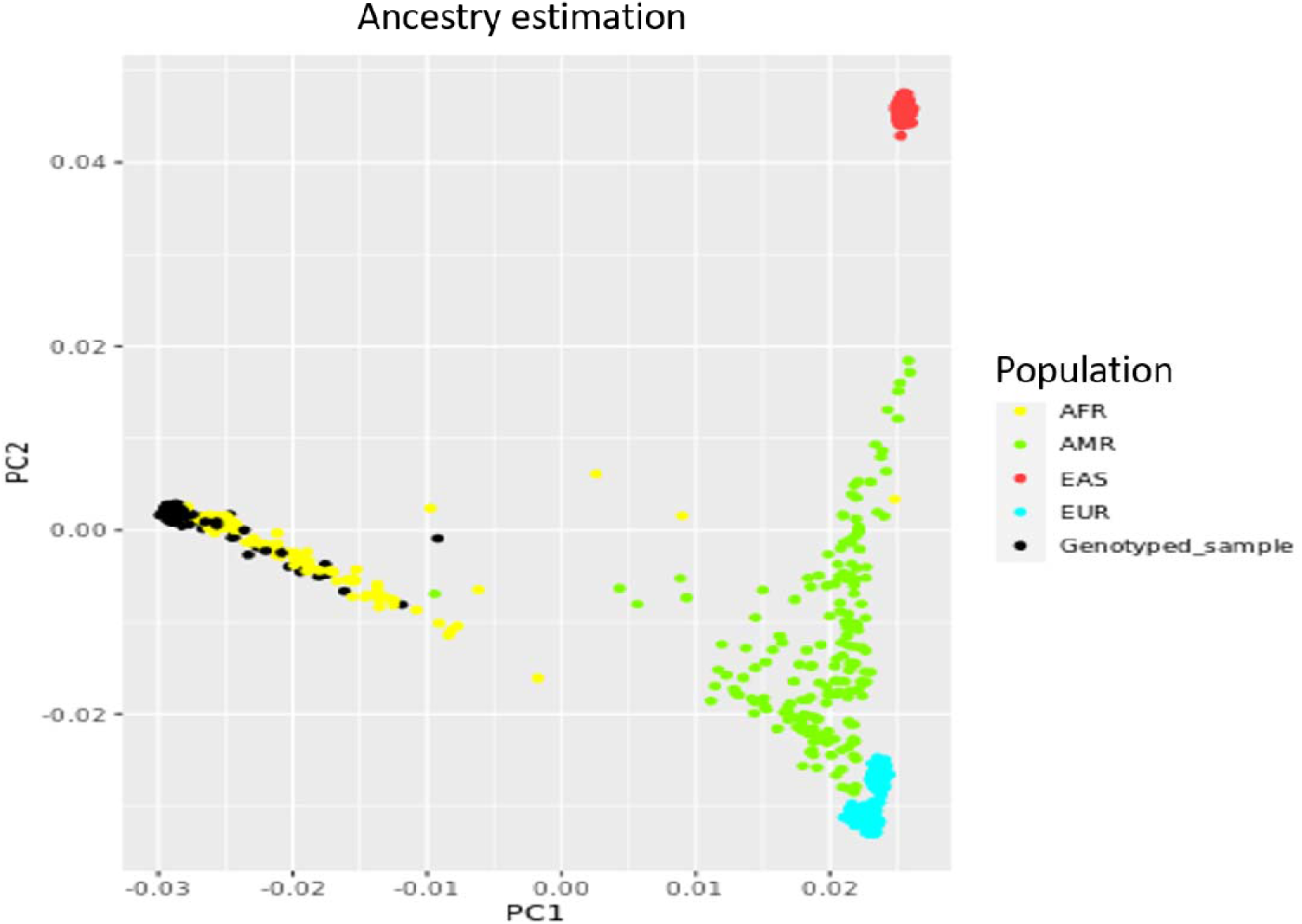
Ancestry estimation in the sickle cell cohort. COPILOT merges study data with the 1000 genome, appropriately prunes the data, performs PCA and plots principal components 1 (PC1) and 2 (PC2). Each dot represents a sample, with colours indicating ancestry based on the 1000 genome. The black dots are representing the study samples. COPILOT correctly predicts that the samples from the sickle cell cohort are all of African descent. Ethnicities are as follows AFR: African, AMR: Ad Mixed American, EAS: East Asian, EUR: European.

**Figure 7:**
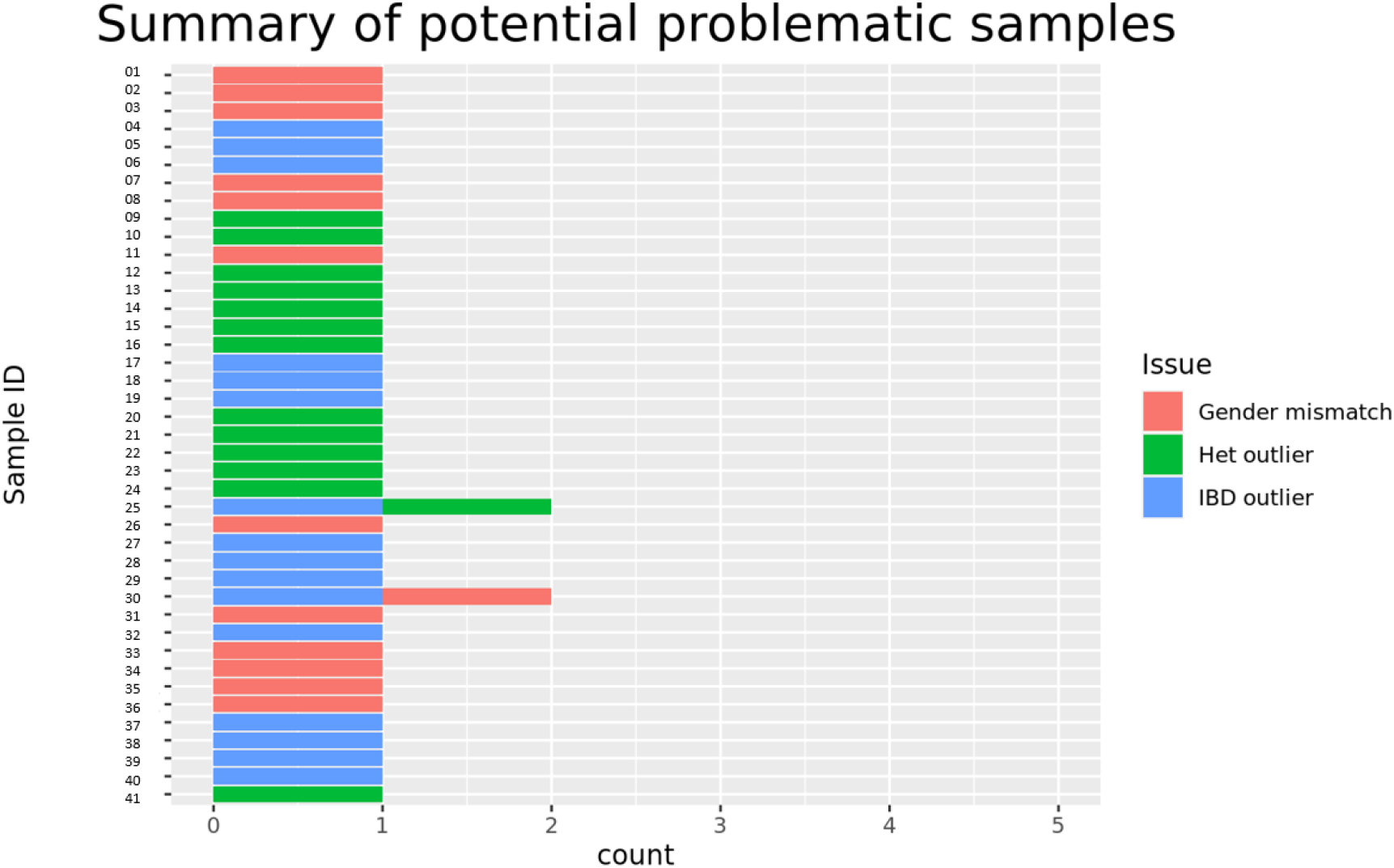
Summary of potential problematic samples in the sickle cell cohort. COPILOT produces a summary of potential outliers and identifies sample outliers dues to multiple criteria, which is often an indication of poor sample quality.

## Discussion

Processing raw Illumina microarray genotyping data involves a battery of complex QC and analysis procedures, which can lead to reproducibility errors, be time-consuming, and be a daunting task for bioinformaticians. The COPILOT protocol addresses these issues by providing an in-depth and clear guide to process raw Illumina genotype data in GenomeStudio, followed by a containerised workflow to automate an array of complex bioinformatics analyses to further process, improve quality and help the user understand the data through an informative and interactive summary report.

The COPILOT protocol was demonstrated on two independent cohorts and showed its ability to improve data quality through significant improvement in sample call rates across all samples processed. More importantly, following the COPILOT protocol rescued 141 samples from a total of 3270 samples, where call rates were below 98% prior to QC and which would have ordinarily been removed from a typical GWAS. Furthermore, as demonstrated in the sickle cell cohort, the COPILOT container automated the identification of samples that require addressing in a GWAS and generated clear and informative visual aids to illustrate the potential issues for the user. The COPILOT protocol is currently deployed in the KING’S College London’s (KCL) Institute of Psychiatry, Psychology and Neuroscience’s (IoPPN) Genomics & Biomarker Core Facility, and has been successfully used to process thousands of samples on various genotyping beadarrays and has proved to be a valuable tool to guide the end data user.

## Conclusions

COPILOT makes processing Illumina genotyping data effortless, reproducible, deployable on multiple platforms, significantly improves sample call rates and reduces the time spent on the data QC process. COPILOT is actively being developed to incorporate additional features, including imputation.

## Availability and requirements

- Project name: COPILOT (a Containerised wOrkflow for Processing ILlumina genOtyping data)
- Project home page: https://khp-informatics.github.io/COPILOT/
- Operating system(s): Platform independent
- Programming language: N/A
- Other requirements: Docker
- Any restrictions to use by non-academics: None
- Contact

## Declarations

### Ethics approval and consent to participate

N/A

### Consent for publication

N/A

### Availability of data and materials

The datasets used and analysed during the current study are available from the corresponding author on reasonable request.

## Funding and acknowledgements

This study presents independent research supported by the NIHR BioResource Centre Maudsley at South London and Maudsley NHS Foundation Trust (SLaM) & Institute of Psychiatry, Psychology and Neuroscience (IoPPN), King’s College London. The views expressed are those of the author(s) and not necessarily those of the NHS, NIHR, Department of Health or King’s College London. RJBD is supported by 1. Health Data Research UK, which is funded by the UK Medical Research Council, Engineering and Physical Sciences Research Council, Economic and Social Research Council, Department of Health and Social Care (England), Chief Scientist Office of the Scottish Government Health and Social Care Directorates, Health and Social Care Research and Development Division (Welsh Government), Public Health Agency (Northern Ireland), British Heart Foundation and Wellcome Trust. 2. The National Institute for Health Research University College London Hospitals Biomedical Research Centre. SM and OO, vas well as the array genotyping of the sickle cell patient group, are supported by MRC grant MR/T013389/1

## Authors’ contributions

HP and RJBD conceptualised the work. HP and SHL developed the COPILOT GenomeStudio SOP. HP designed and developed the COPILOT container. HP created the online documentations and COPILOT website. SM and OO generated and processed the sickle cell cohort. GB and SHL generated and processed the mental health cohort. All authors contributed and reviewed the final manuscript.

